# HOS1 promotes plant tolerance to low-energy stress *via* the SnRK1 protein kinase

**DOI:** 10.1101/2023.01.26.525711

**Authors:** Leonor Margalha, Alexandre Elias, Borja Belda-Palazón, Bruno Peixoto, Ana Confraria, Elena Baena-González

**Author notes:** Equal contribution (Leonor Margalha, Alexandre Elias). Department of Biology, University of Oxford, South Parks Rd, Oxford, OX1 3RB, UK.

## Abstract

Plants need to integrate internal and environmental signals to mount adequate stress responses. The NUCLEAR PORE COMPLEX (NPC) component HIGH EXPRESSION OF OSMOTICALLY RESPONSIVE GENES 1 (HOS1) is emerging as such an integrator, affecting responses to cold, heat, light and salinity. Stress conditions often converge in a low-energy signal that activates SUCROSE NON-FERMENTING 1-RELATED KINASE 1 (SnRK1) to promote stress tolerance and survival. Here, we explored the role of HOS1 in the SnRK1-dependent response to low-energy stress in *Arabidopsis thaliana*, using darkness as a treatment and a combination of genetic, biochemical and phenotypic assays. We show that the induction of starvation genes and plant tolerance to prolonged darkness are defective in the *hos1* mutant. HOS1 interacts physically with the SnRK1α1 catalytic subunit in yeast-two-hybrid and *in planta*, and the nuclear accumulation of SnRK1α1 is reduced in the *hos1* mutant. Likewise, another NPC mutant, *nup160*, exhibits lower activation of starvation genes and decreased tolerance to prolonged darkness. Importantly, defects in low-energy responses in the *hos1* background are rescued by fusing SnRK1α1 to a potent nuclear localization signal, or by sugar supplementation during the dark treatment. Altogether, this work demonstrates the importance of HOS1 for the nuclear accumulation of SnRK1α1, which is key for plant tolerance to low-energy conditions.

## INTRODUCTION

Abiotic environmental stresses such as high and low temperature, drought, flooding, and salinity can be yield-limiting factors that compromise crop productivity. To cope with such adverse environmental conditions, plants have developed responses that promote stress tolerance and survival, usually at the expense of growth (Margalha *et al*., 2019).

The *HIGH EXPRESSION OF OSMOTICALLY RESPONSIVE GENE 1* (*HOS1*) locus was initially identified as a negative regulator of cold signalling in Arabidopsis (Ishitani *et al*., 1998; Lee *et al*., 2001; Dong *et al*., 2006). Subsequent studies unveiled the involvement of HOS1 in the response to other environmental signals, such as high temperature (Zhang *et al*., 2020; Han *et al*., 2020), light quality (MacGregor *et al*., 2013; Lazaro *et al*., 2015) and salinity (Zhu *et al*., 2017). HOS1 has been implicated in auxin (Lee and Seo, 2015a), ethylene (Lee and Seo, 2015b) and ABA (Zhu *et al*., 2017), as well as in developmental processes such as germination (Lazaro *et al*., 2015), primary root growth (Zhu *et al*., 2017), hypocotyl elongation (MacGregor *et al*., 2013; Lee and Seo, 2015a; Kim, Lee, Jung, *et al*., 2017; Kim, Lee and Park, 2017; Zhang *et al*., 2020), leaf expansion (Lee and Seo, 2015b; Zhang *et al*., 2020) and flowering (Lazaro *et al*., 2012; Lee *et al*., 2012; Jung *et al*., 2012; Jung *et al*., 2013; Seo *et al*., 2013; Lazaro *et al*., 2015).

HOS1 harbors a RING finger domain near the N-terminus of the protein (Lee *et al*.,2001) and acts as an E3 ubiquitin ligase promoting the proteasome-dependent degradation of the transcription factors INDUCER OF CBF EXPRESSION 1 (ICE1) (Dong *et al*.,2006) and CONSTANS (CO) (Lazaro *et al*., 2012), involved in cold acclimation and in the photoperiodic regulation of flowering, respectively. HOS1 also functions as a chromatin modifier, contributing to the transcriptional activation of the flowering repressor gene *FLOWERING LOCUS C (FLC*) (Jung *et al*., 2013), the microRNA gene *MIR168b*, involved in *ARGONAUTE 1 (AGO1*) regulation (Wang *et al*., 2015), and DNA repair genes such as *RECQ2*, encoding a DNA helicase important for thermotolerance (Han *et al*., 2020). Such functions are likely to be performed through association with chromatin binding factors since a direct association to DNA has not been demonstrated. In these cases, as well as in the case of the transcription factor PHYTOCHROME INTERACTING FACTOR 4 (PIF4), HOS1 modulates protein function through mechanisms that do not involve proteasomal degradation (Lee *et al*., 2012; Kim, Lee, Jung, *et al*., 2017; Han *et al*., 2020; Jung *et al*., 2013). On the other hand, HOS1 is the only protein in the Arabidopsis genome with a region of homology to the yeast and animal nucleoporin (NUP) ELYS (Jung *et al*., 2014). HOS1 interacts directly with the nuclear pore complex (NPC) components NUP96 (Cheng *et al*.,2020) and NUP160 (Li *et al*., 2020), and co-immunopurifies with RAE1, NUP43 (Tamura *et al*., 2011), and NUP85 (Zhu *et al*., 2017). With the exception of RAE1, all the mentioned NUPs belong to the NUP107-160 subcomplex, the major constituent of the outer ring complex of the NPC core scaffold, with essential structural functions (Li and Gu, 2020). These interactions support a role for HOS1 in the NPC-mediated control of nucleo-cytoplasmic trafficking. In line with such role, *hos1* plants show nuclear retention of poly-adenylated mRNAs, which may underlie the lengthened circadian period observed in this mutant (MacGregor *et al*., 2013). In addition, nuclear accumulation of the PIF4 transcription factor is reduced in *hos1*, compromising the response of the mutant to warm temperatures (Zhang *et al*., 2020). Several other *nup* mutants are, like *hos1*, also affected in mRNA export and nucleo-cytoplasmic trafficking of proteins, temperature signalling, circadian function, plant growth and flowering time (Zhang *et al*., 2020; MacGregor and Penfield, 2015).

Most types of environmental stress compromise photosynthesis or/and respiration (Muhammad *et al*., 2021; Cho *et al*., 2021), thereby leading to reduced ATP production and low-energy stress (Branco-Price et al., 2008; de Col et al., 2017; Tomé et al., 2014). One major player in the sensing of energy resources and the orchestration of adequate downstream responses is the evolutionarily conserved SUCROSE NON-FERMENTING 1 (SNF1)-RELATED KINASE 1 (SnRK1), the plant ortholog of mammalian AMP-ACTIVATED KINASE (AMPK) and yeast SNF1. SnRK1 is activated when energy levels decline and is repressed by sugars such as trehalose 6-phosphate, glucose 6-phosphate and glucose 1-phosphate (Zhang *et al*., 2009; Nunes *et al*., 2013; Zhai *et al*., 2018; Peixoto *et al*., 2021). Upon activation, SnRK1 drives vast transcriptional and metabolic adjustments to stimulate energy-producing catabolic processes and inhibit energy-consuming biosynthetic processes and growth (Baena-González *et al*., 2007; Baena-González and Sheen, 2008; Cho *et al*., 2012; Pedrotti *et al*., 2018). Constitutive manipulation of SnRK1 levels results in altered tolerance to a wide range of stresses (Hulsmans *et al*., 2016; Margalha *et al*., 2019), and affects multiple aspects of growth and development (Baena-González and Hanson, 2017; Jamsheer K *et al*., 2021).

Like its mammalian and yeast orthologs, SnRK1 functions as a heterotrimeric complex composed of an α-catalytic, and β- and γ-regulatory subunits, encoded by different genes (*SnRK1α1/α2, SnRK1β1/β2/β3*, and *SnRK1βγ*, in Arabidopsis). Despite the functional and structural conservation, SnRK1 displays unique features that may underlie functions and modes of regulation specific to plants (Ramon *et al*., 2013; Emanuelle *et al*., 2015; Broeckx *et al*., 2016). Recent studies have highlighted the importance of subcellular localization for SnRK1 function, with clear phenotypes associated to the confinement of SnRK1 inside or outside of the nucleus (Ramon *et al*., 2019; Gutierrez-Beltran and Crespo, 2022). The nuclear localization of SnRK1 is essential for its role in transcriptional regulation and is promoted by stresses that cause an energy deficit, although this response may be limited to specific cell types or tissues (Cho *et al*., 2012; Ramon *et al*., 2019; Blanco *et al*., 2019). The nuclear localization of SnRK1 responds also to N (Yuan *et al*., 2016; Sun *et al*., 2021) and to ABA signalling (Belda-Palazón *et al*., 2022), and displays a diurnal oscillation pattern (Yuan *et al*., 2016). On the other hand, SnRK1 phosphorylates metabolic enzymes [e.g. SUCROSE PHOSPHASE SYNTHASE (SPS) and NITRATE REDUCTASE (NR) (Sugden *et al*., 1999; Nukarinen *et al*., 2016)] that reside outside the nucleus. SnRK1 was also found to associate with the ER (Blanco *et al*., 2019) where it interacts with FCS-like zinc finger (FLZ) proteins (Jamsheer K *et al*., 2018), and to the ER perinuclear ring (Blanco *et al*., 2019), where it co-localizes with the component of the TARGET OF RAPAMYCIN (TOR) complex, REGULATORY ASSOCIATED PROTEIN OF TOR 1B (RAPTOR1B) (Nukarinen *et al*.,2016). However, the mechanisms that regulate the subcellular distribution of SnRK1 are poorly understood.

In this study we show that HOS1 is important for plant tolerance to low-energy conditions and that this is largely due to its involvement in the accumulation of SnRK1α1 in the nucleus.

## RESULTS

### HOS1 is required for low-energy responses

HOS1 is a multifaceted protein with a central role in diverse stress responses (MacGregor and Penfield, 2015). Given that numerous stress conditions lead to low-energy levels (Baena-González and Sheen, 2008; Tomé et al., 2014), we wondered whether HOS1 could be involved in the response to low-energy stress. To investigate this, we compared the induction of starvation genes by reverse transcription quantitative PCR (RT-qPCR) in leaves of wild-type Col-0 and the *hos1-3* knockout mutant [hereafter referred as *hos1;* (Lazaro *et al*., 2012)] in response to a 3 h dark treatment. Starvation genes are involved in metabolic rearrangements and nutrient remobilization strategies and are essential for coping with low-energy stress (Baena-González *et al*., 2007; Baena-González and Sheen, 2008; Cookson *et al*., 2016; Ramon *et al*., 2019; Pedrotti *et al*., 2018). As shown in Fig. **S1A**, a 3 h dark treatment was sufficient to activate starvation genes such as *DIN6, DIN1* and *DRM2* in Col-0 but the induction of these genes was reduced in the *hos1* mutant. These defects were partially rescued in a *35S::HOS1-GFP/hos1* line *13-9* (hereafter referred as *HOS1-GFP*, Fig. **S1A,D**). These results suggested the involvement of HOS1 in the response to low-energy stress.

Successful activation of starvation genes is associated with survival after prolonged periods of darkness (Pedrotti *et al*., 2018). We therefore tested whether the inability of *hos1* to mount an adequate starvation response at the molecular level led to decreased plant tolerance to a prolonged dark stress. To this end, we transferred 4-week-old plants to darkness for 7 days (d) and placed them back to light for 7 d to recover. In agreement with the molecular response from 3h dark treatments, *hos1* showed decreased plant tolerance to prolonged darkness, manifested as yellowing and wilting of the leaves at 7 d (Fig. **S1B**), and as compromised growth or complete growth arrest during the subsequent recovery period (7+ 7 d) (Fig. **S1B**). The differences in the sensitivity to prolonged darkness between Col-0 and *hos1* were also reflected in the gain in rosette fresh weight determined at 7 d and at 7+ 7 d as compared to day 0 (Fig. **S1C**). Furthermore, the decreased plant tolerance of *hos1* was partially reverted in the *HOS1-GFP* complementation line (Fig. **S1B-C**). A different mutant allele of *HOS1, hos1-4* (Lazaro *et al*., 2012), was also analyzed (Fig. **S2**). To better monitor the molecular response to a prolonged dark treatment, samples were collected 2 d after the onset of darkness. Similarly to *hos1-3* (Fig. **S1**), *hos1-4* plants showed defective induction of starvation genes (Fig. **S2A**) and decreased plant tolerance (Fig. **S2B**), with a lower gain in rosette fresh weight than Col-0 after the dark treatment (Fig. **S2C**). Altogether, these results support a role for HOS1 in plant tolerance to prolonged darkness.

Starvation genes are under the control of the SnRK1 protein kinase that is activated in response to low-energy conditions and repressed by sugars (Baena-González *et al*., 2007; Baena-González and Sheen, 2008; Pedrotti *et al*., 2018; Ramon *et al*., 2019). SnRK1 is also required for surviving prolonged periods of darkness (Pedrotti *et al*., 2018; Ramon *et al*., 2019; Henninger *et al*., 2022). To investigate if the defects of *hos1* in low-energy signalling are SnRK1-dependent, we crossed *hos1* to a *SnRK1α1* loss-of-function mutant, *snrk1α1-3* [hereafter referred as *snrk1α1;* (Mair *et al*., 2015; Crozet *et al*., 2016)], and assessed the response of the double mutant to darkness. The 2 d dark treatment led to defective induction of starvation genes in *hos1* (Fig. **1A**) and in the *snrk1α1* mutant, confirming previous reports that this response is SnRK1-dependent (Pedrotti *et al*., 2018; Ramon *et al*., 2019; Henninger *et al*., 2022). However, *snrk1α1* plants displayed only a mild sensitivity to prolonged darkness (Fig. **1B-C**). This is in agreement with the functional redundancy between the two α-catalytic subunits of SnRK1, encoded by the *SnRK1α1* and *SnRK1α2* genes in Arabidopsis (Baena-González *et al*., 2007; Ramon *et al*., 2019). The *hos1/snrk1α1* double mutant showed defects in the induction of starvation genes that were either similar to the *snrk1α1* and *hos1* parents or slightly enhanced (Fig. **1A**). The sensitivity of *hos1/snrk1α1* to prolonged darkness was similar to that of *hos1*, both being severely compromised or unable to resume growth during the 7 d recovery period (Fig. **1B-C**). Overall, these results suggest HOS1 is required for adequate SnRK1 signalling in response to low-energy stress.

**Figure 1.**
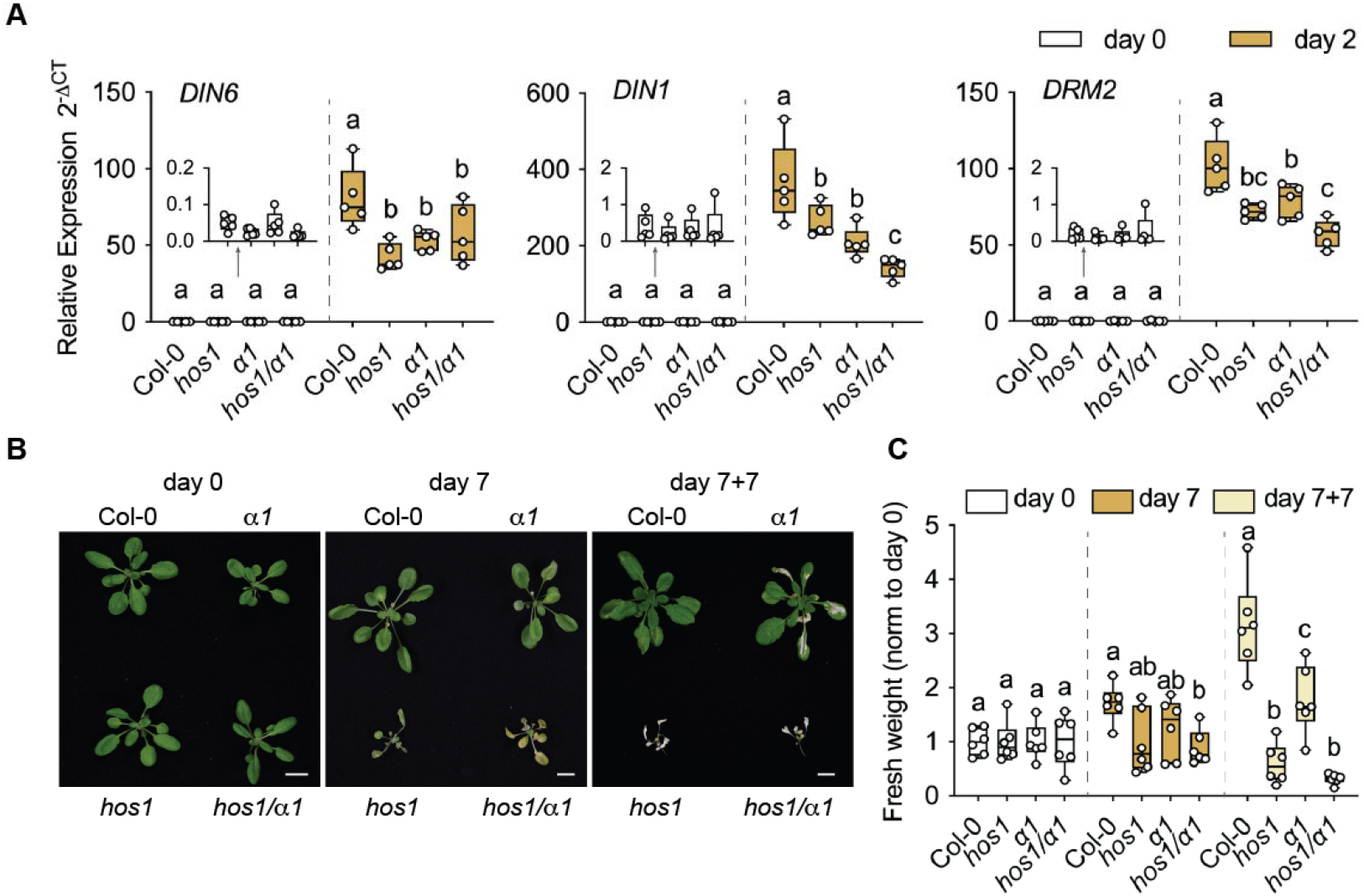
Tolerance of *hos1* and *Snrk1α1* mutants to prolonged darkness. **A.** Induction of starvation genes (*DIN6, DIN1, DRM2*) measured by RT-qPCR from rosettes of Col-0, *α1* (*snrk1α1*), *hos1*, and *hos1/α1* plants collected at day 0 and after 2 d of dark treatment (day 2). Box and whiskers plots represent 5 biological replicates (each consisting of 1 rosette per genotype and condition, grown in 5 independent batches). **B.** Representative pictures of 4-week-old plants (day 0) of the indicated genotypes placed in constant darkness for 7 days (day 7) and transferred back to the initial photoperiod for 7 additional days (day 7+7). Scale bar = 1 cm). **C.** Rosette fresh weight determined at days 0, 7 and 7+7, and normalized to day 0 for each genotype. Box and whiskers plots represent 6 plants per genotype and condition (grown in 3 independent batches). Different letters denote statistically significant differences between genotypes within each condition (days 0, 2, 7 and 7+7), as determined by two-way ANOVA with Tukey’s multiple comparisons test (p<0.05).

To further determine if *hos1* sensitivity to prolonged darkness resulted indeed from the ensuing energy deprivation, Col-0 and *hos1* were grown in 0.5 x MS media plates with or without 1% sucrose and subjected to a prolonged dark treatment (Fig. **S3**). In agreement with a role for HOS1 in the low-energy stress response, sugar supplementation could largely rescue the sensitive phenotype of *hos1* at 7+ 7 d and increase the survival rate of *hos1* seedlings from 29% in control media to 100% in the presence of sugar (Fig. **S3A-B**).

### Nuclear accumulation of SnRK1α1 requires HOS1

We next performed protein-protein interaction studies to investigate whether the requirement of HOS1 for proper SnRK1 signalling could be based on a physical interaction between HOS1 and SnRK1α1 (Fig. **2**). In yeast two-hybrid (Y2H) assays, cell growth in highly stringent media was only observed when both proteins were co-expressed (Fig. **2A**), indicating a direct interaction between SnRK1α1 and HOS1 in yeast cells. To assess if SnRK1α1 interacts with HOS1 also *in planta*, we performed co-immunoprecipitation (co-IP) experiments, pulling down GFP from *HOS1-GFP* or *GFP* control plants. Subsequent Western-blot (WB) detection with an antibody against SnRK1α1, showed that SnRK1α1 co-purified with HOS1-GFP but not with GFP alone (Fig. **2B**).

**Figure 2.**
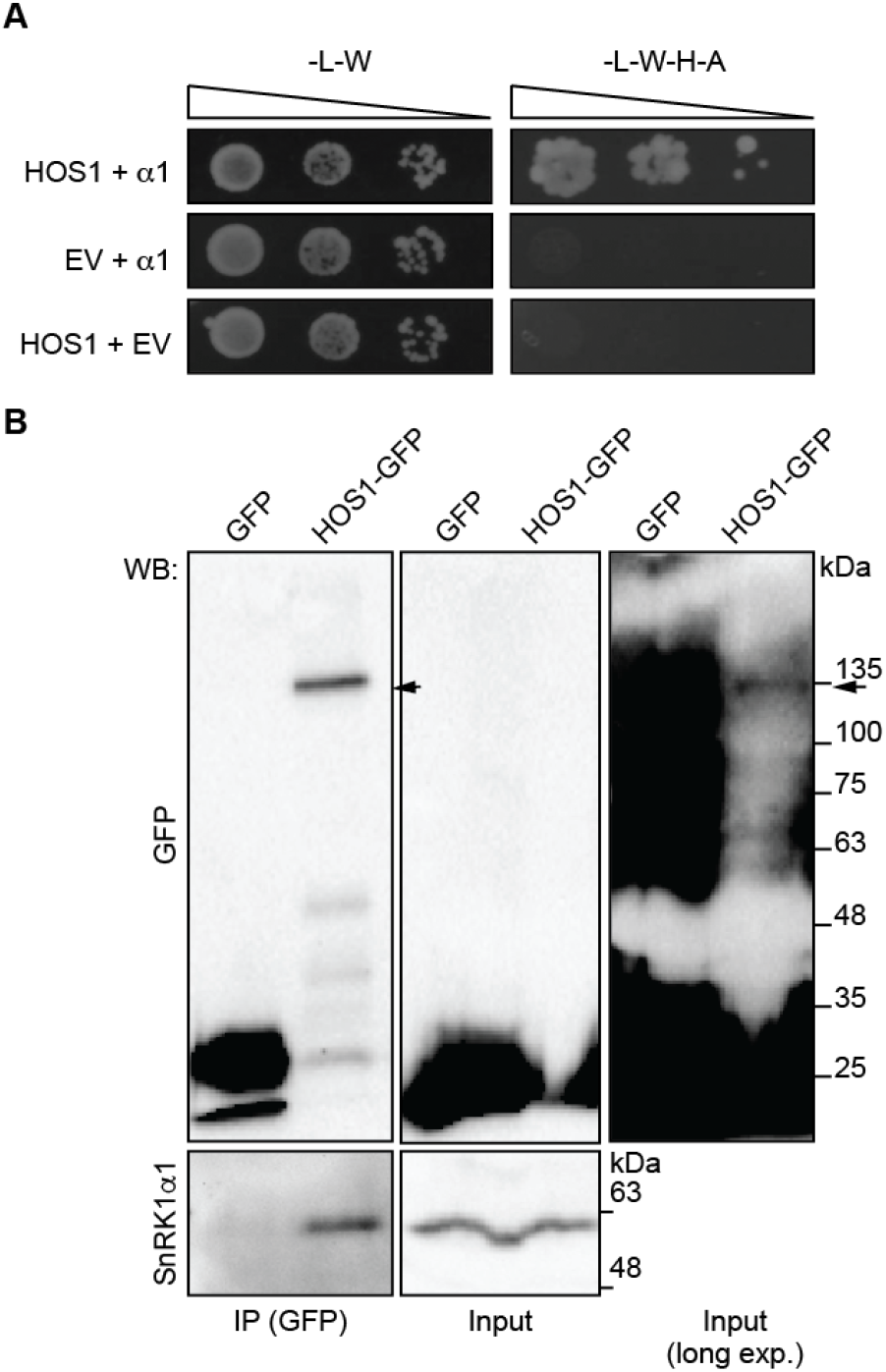
Physical interaction between HOS1 and SnRK1□1. **A.** Yeast two-hybrid assays. Yeast cell growth after expression of HOS1 and SnRK1α1 (α1), together or alone with the complementary empty vectors (EV), in low (-L, -W) or high (-L, -W, -H, -A) stringent media. (L) leucine; (W) tryptophan; (H) histidine; (A) adenine. Representative image of 3 biological replicates, grown in 3 independent batches. **B.** GFP immunoprecipitation (IP) from rosettes of *35S::GFP (GFP*) or *35S::HOS1-GFP/hos1 (HOS1-GFP*) 4-week-old plants. Western-blot (WB) with antibodies against GFP and SnRK1α1. Right panel, longer exposure of the middle panel to visualize HOS1-GFP in the input. Black arrows point to HOS1-GFP. Representative image of 3 biological replicates (each consisting of 1 rosette per genotype, grown in 3 independent batches).

Given the physical interaction between HOS1 and SnRK1α1, we next asked if HOS1 could affect SnRK1 stability. HOS1 has an E3 ubiquitin ligase activity that is key for many environmental and developmental responses (Dong *et al*., 2006; Lee *et al*., 2012; Jung *et al*., 2012; Lazaro *et al*., 2012; Lazaro *et al*., 2015). Furthermore, HOS1 and the SUMO E3 ligase SIZ1 act antagonistically on ICE1 stability (Miura *et al*., 2007; Miura *et al*., 2011), and SIZ1 has been implicated in the SUMO-dependent proteasomal degradation of SnRK1 (Crozet *et al*., 2016). However, no clear differences were observed in total protein extracts between Col-0 and *hos1* regarding the accumulation of the SnRK1α1 and SnRK1α2 catalytic subunits, their T-loop phosphorylation, necessary for kinase activity (Baena-González *et al*., 2007; Shen *et al*., 2009), or the levels of the SnRK1β1 regulatory subunit (Fig. **S4**). These results suggest that the increased sensitivity of *hos1* in response to prolonged darkness is unrelated to changes in total SnRK1 protein accumulation.

HOS1 localization at the NPC is essential for some of its functions (Li *et al*., 2020). On the other hand, the activation of starvation genes by SnRK1 requires its presence in the nucleus (Cho *et al*., 2012; Ramon *et al*., 2019). Therefore, we asked whether HOS1 could affect specifically the nuclear pool of SnRK1α1. To test this, we performed nuclear fractionation from Col-0 and *hos1* rosettes and analyzed SnRK1α1 in the resulting fractions (Fig. **3**). No significant differences were observed in SnRK1α1 levels in the cytoplasmic fractions of Col-0 and *hos1* (Fig. **3A-B**). However, the levels of SnRK1α1 protein were reduced in the nuclear fraction of *hos1* in comparison to Col-0 (Fig. **3A-B**). A lower accumulation of SnRK1α1 in the nucleus of *hos1* was also observed after a 9h dark treatment (Fig. **3C-D**), reported to induce nuclear translocation of SnRK1α1 (Ramon *et al*., 2019), indicating that HOS1 is important for the accumulation of SnRK1α1 in the nucleus both under control and low-energy conditions.

**Figure 3.**
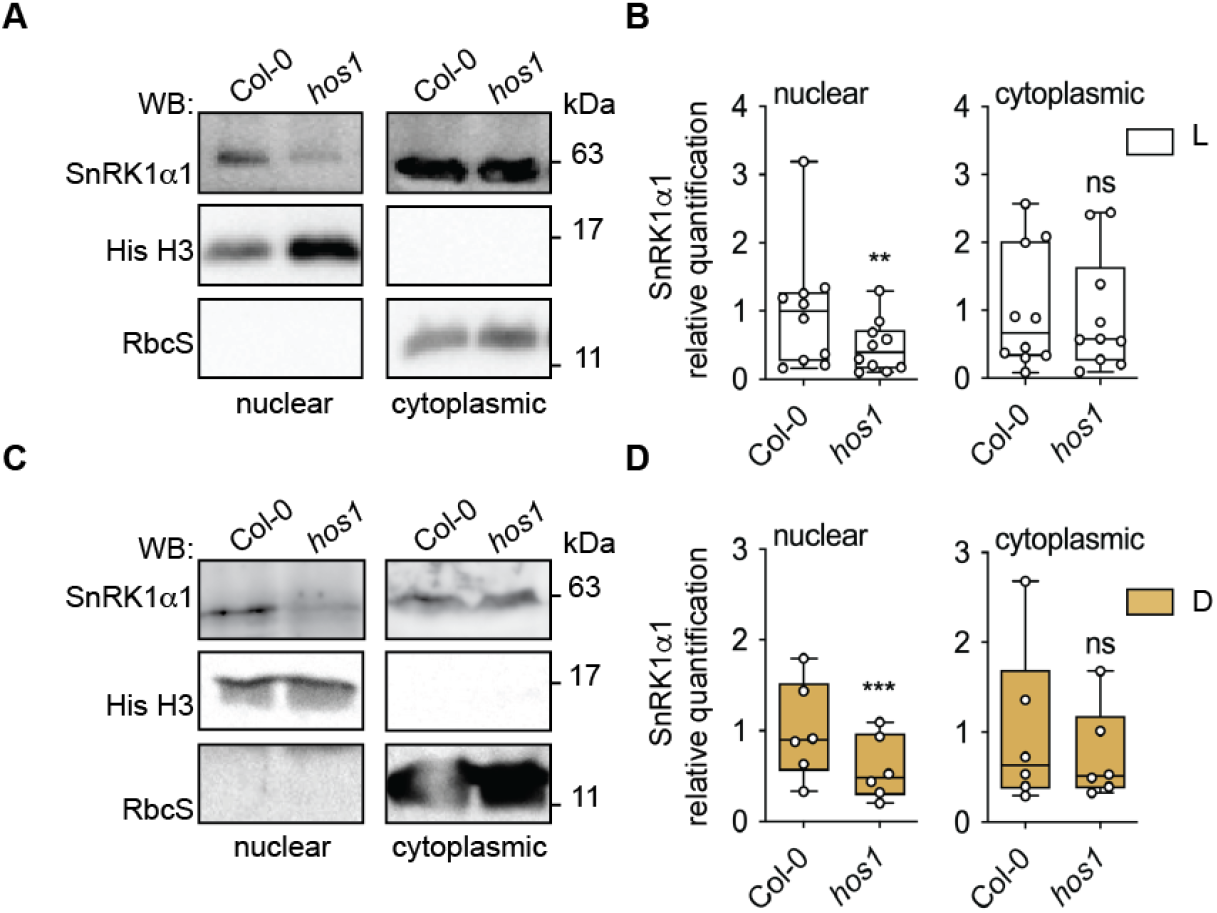
Nuclear accumulation of SnRK1α1 in *hos1*. **A-D.** Nuclear fractionation from rosettes of 4-week-old Col-0 and *hos1* plants under control light (L) conditions (**A-B**) or after a 9 h night extension (dark, D; **C-D**). **A,C**. Representative Western-blots (WB) of nuclear and cytoplasmic fractions with antibodies against SnRK1α1, Histone 3 (His H3, nuclear marker, not detected in the WB image of the cytoplasmic fraction), and Rubisco small subunit (RbcS, cytosolic marker, not detected in the WB image of the nuclear fraction). **B,D.** SnRK1α1 quantification normalized to His H3 in the nuclear fraction and to RbcS in the cytoplasmic fraction, and further normalized to Col-0. Box and whiskers plots represent a minimum of 6 biological replicates (each consisting of 1 rosette per genotype and condition, grown in a minimum of 6 independent batches). Asterisks denote statistically significant differences between Col-0 and *hos1*, as determined by a ratio paired *t*-test. (**) p< 0.01; (***) p< 0.001; (*ns*), non-significant.

These data suggested a constitutive effect of HOS1 on the nuclear accumulation of SnRK1α1. To investigate if this effect is associated with the NPC localization or function of HOS1, we examined the response to low-energy stress in the *nup160* mutant (Fig. **4**) since the NUP160 subunit is required for anchoring HOS1 at the NPC (Li *et al*., 2020). Similarly to *hos1, nup160* showed defective activation of starvation genes in response to 2 d of darkness (Fig. **4A**), decreased plant tolerance to an extended 7 d period of darkness and a compromised ability to resume growth in the subsequent recovery period (Fig. **4B-C**). This suggests that anchoring HOS1 at the NPC, or NPC integrity, is important for nuclear accumulation of SnRK1α1 and for an adequate starvation response. Moreover, a double *nup160/hos1* mutant showed similar plant phenotypes to prolonged darkness as the single parents (Fig. **S5**), supporting that the two nucleoporins act in the same pathway to promote tolerance to low-energy stress.

**Figure 4.**
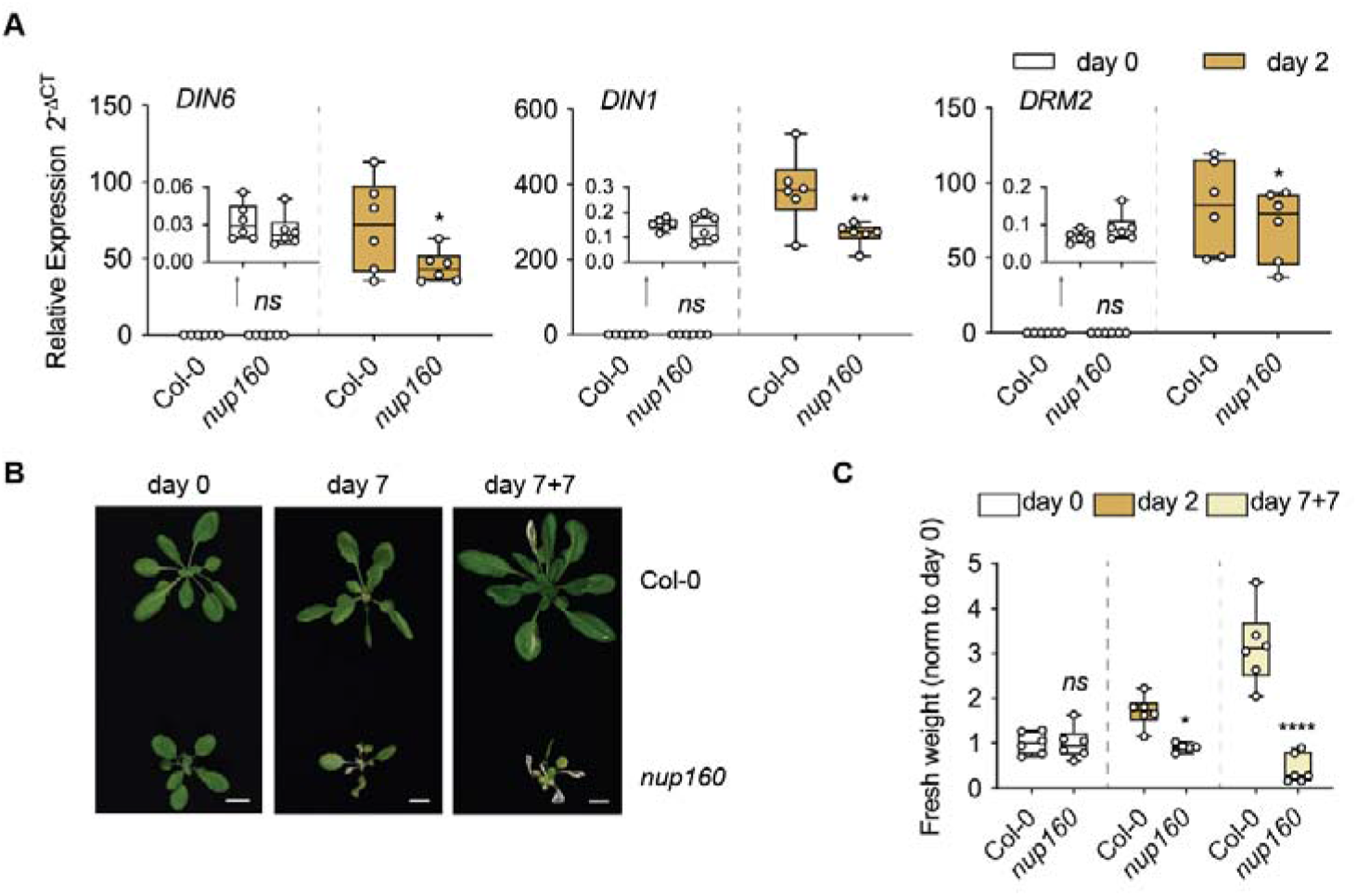
Tolerance of the NPC mutant *nup160* to prolonged darkness. **A.** Induction of starvation genes (*DIN6, DIN1, DRM2*) measured by RT-qPCR from rosettes of 4-week-old Col-0 and *nup160* mutant plants collected at day 0 and after 2 d of dark treatment (day 2). Box and whiskers plots represent 6 biological replicates (each consisting of 1 rosette per genotype and condition, grown in 6 independent batches). **B.** Representative pictures of 4-week-old Col-0 and *nup160* plants (day 0) placed in constant darkness for 7 days (day 7) and transferred back to the initial photoperiod for 7 additional days (day 7+7). Scale bar = 1 cm. **C.** Rosette fresh weight determined at days 0, 7 and 7+7, and normalized to day 0 for each genotype. Box and whiskers plots represent 5-6 plants per genotype and condition (grown as 3 independent batches; note that the Col-0 set is the same as in Fig. **1C**). Asterisks denote statistically significant differences between Col-0 and *nup160* within each condition (days 0, 2, 7 and 7+7), as determined by two-way ANOVA with Sidak’s multiple comparisons test. (*) p< 0.05; (**) p< 0.01; (****) p< 0.0001; (*ns*), non-significant.

### Fusion of SnRK1α1 to a potent nuclear localization signal (NLS) signal rescues *hos1* defects in low-energy responses

We next wondered if depletion of SnRK1α1 from the nucleus (Fig. **3A-D**) was sufficient to explain the molecular and plant phenotypes of *hos1* in response to darkness (Fig. **1**, Fig. **S1**, Fig. **S3** and Fig. **S5**). To assess this, we crossed the *hos1* mutant to two recently described lines, in which the *snrk1α1/snrk1α2* double mutant was complemented with SnRK1α1 variants with different subcellular localizations (Ramon *et al*., 2019). In one line, the simian virus 40 large T-antigen NLS (SV40-NLS) was fused *N*-terminally to SnRK1α1 to increase its nuclear localization [Fig. **5**, *NLS-α1 (NLS-SnRK1α1/snrk1α1/snrk1α2)]*. In the other line, the N-terminal myristoylation motif of the SnRK1β2-subunit was fused *N*-terminally to SnRK1α1 to increase its membrane association and exclusion from the nucleus [Fig. **5**, *βMYR-α1* (*βMYR-SnRK1α1/snrk1α1/snrk1α2*)]. Nuclear fractionation from *hos1* and *hos1/NLS-α1 (NLS-SnRK1α1/snrk1α1/snrk1α2/hos1*) plants showed that fusing the SV40-NLS motif to SnRK1α1 can indeed increase its nuclear accumulation in the *hos1* background (Fig. **5A-B**). Most importantly, the *NLS-SnRK1α1* transgene in the *hos1/NLS-α1* plants was able to restore normal induction of starvation genes in response to darkness (Fig. **5C**) and rescue to a large extent the *hos1* sensitivity to the 7 d dark treatment and recovery (Fig. **5D**), with a significant increase in the fresh weight gain in the *hos1/NLS-α1* line in comparison to *hos1* (Fig. **5E**). By contrast, the *βMYR-SnRK1α1* transgene in *hos1/βMYR-α1* plants (*βMYR-SnRK1α1/snrk1α1/snrk1α2/hos1*) aggravated the defects of *hos1* in the induction of starvation genes in response to darkness (Fig. **5C**). Consequently, expression of *βMYR-SnRK1α1* exacerbated the sensitivity of *hos1* to a prolonged dark treatment (Fig. **5D-E**). Collectively, these results suggest that the molecular and plant phenotypes of *hos1* in response to low-energy conditions are at least partially caused by the reduced nuclear accumulation of SnRK1α1.

**Figure 5.**
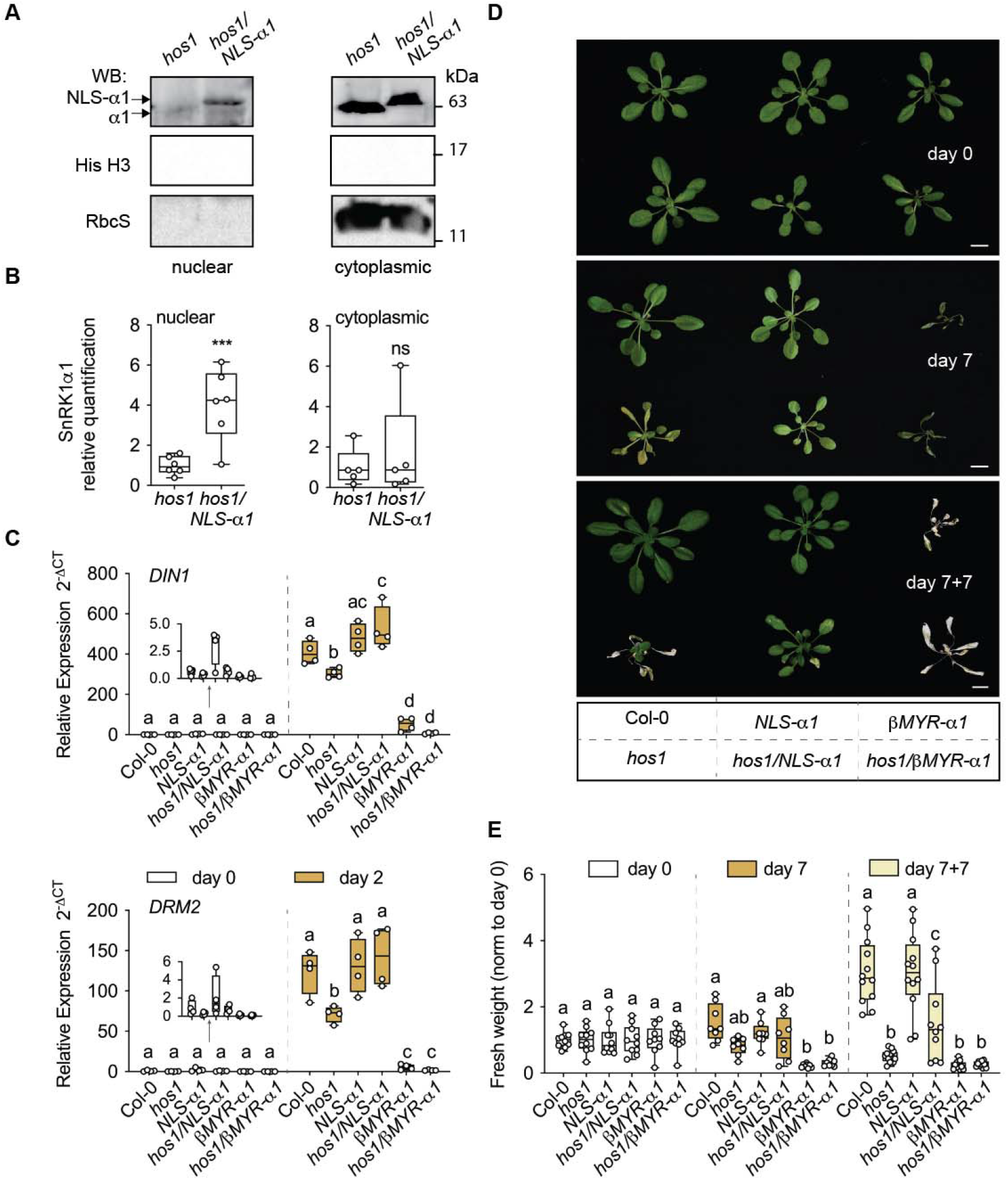
Impact of *NLS-SnRK1α1* in *hos1* tolerance to prolonged darkness. Cross between *hos1* and plants with predominant localization of SnRK1α1 in the nucleus (*NLS-SnRK1α1*, shown as *NLS-α1*) or outside the nucleus (*βMYR-SnRK1α1*. shown as *βMYR-α1*) (Ramon *et al*., 2019). **A.** Nuclear fractionation from rosettes of 4-week-old *hos1* and *hos1/NLS-α1* plants. Representative Western-blots (WB) of nuclear and cytoplasmic fractions with specific antibodies against SnRK1α1, Histone 3 (His H3, nuclear marker, not detected in the WB image of the cytoplasmic fraction), and Rubisco small subunit (RbcS, cytosolic marker, not detected in the WB image of the nuclear fraction). Arrows point to SnRK1α1 (α1) or NLS-α1 proteins. **B.** SnRK1α1 quantification normalized to His H3 in the nuclear fraction, to RbcS in the cytoplasmic fraction, and further normalized to *hos1*. Box and whiskers plots represent a minimum of 5 biological replicates, grown in a minimum of 5 independent batches. Asterisks denote statistically significant differences between *hos1* and *hos1/NLS-α1*, determined by a ratio paired *t*-test. (***) p< 0.001; (*ns*), non-significant. **C.** Induction of starvation genes (*DIN1, DRM2*) measured by RT-qPCR, from rosettes of 4-week-old plants of the indicated genotypes collected at day 0 and after 2 d of dark treatment (day 2). Box and whiskers plots represent 4 biological replicates (each consisting of 1 rosette per genotype, grown in 4 independent batches). **D.** Representative pictures of 4-week-old plants (day 0) of the indicated genotypes placed in constant darkness for 7 d (day 7) and transferred back to the initial photoperiod for 7 additional days (day 7+7). Scale bar = 1 cm. **E.** Rosette fresh weight determined at days 0, 7 and 7+7, and normalized to day 0 for each genotype. Box and whiskers plots represent 8-12 plants per genotype and condition (grown in 4 independent batches). Different letters denote statistically significant differences between genotypes within each condition (days 0, 2, 7 and 7+7), as determined by two-way ANOVA with Tukey’s multiple comparisons test (p<0.05).

To investigate the specificity of the rescue provided by the *NLS-SnRK1α1* transgene on *hos1* phenotypes, we also monitored flowering time, since both HOS1 (Lazaro *et al*.,2012; Lee *et al*., 2012; Jung *et al*., 2012; Seo *et al*., 2013; Jung *et al*., 2013) and SnRK1α1 (Tsai and Gazzarrini, 2012; Williams *et al*., 2014; Jeong *et al*., 2015; Sanagi *et al*., 2021; Baena-González *et al*., 2007) were shown to act as negative regulators of this trait. However, the early flowering phenotype of *hos1* in long-day conditions was not modified by changes in SnRK1α1 localization, with both *hos1/NLS-α1* and *hos1/βMYR-α1* plants showing early flowering similar to *hos1* (Fig. **S6**). This indicates that not all *hos1* phenotypes can be complemented by enhancing the levels of SnRK1α1 in the nucleus.

## DISCUSSION

Since its first identification as a negative regulator of cold signalling, the role of HOS1 has expanded towards the integration of environmental and endogenous signals in numerous stress and developmental responses (Ishitani *et al*., 1998; Lee *et al*., 2001; Dong *et al*., 2006; Lazaro *et al*., 2012; Lee *et al*., 2012; MacGregor *et al*., 2013; Lee and Seo, 2015b; Lee and Seo, 2015a; Lazaro *et al*., 2015; Zhu *et al*., 2017; Zhang *et al*., 2020; Han *et al*., 2020). Here, we identify HOS1 as a positive regulator of the low-energy stress response triggered by unexpected or prolonged darkness (Fig. S7).

The *hos1* mutants displayed defective induction of starvation genes (Fig. **S1A**, Fig. **S2A** Fig. **1A**, and Fig. **5C**) and increased sensitivity to prolonged darkness (Fig. **S1B-C**, Fig. **S2B-C**, Fig. **1B-C**, and Fig. **5D-E**). Importantly, sugar supplementation could rescue the *hos1* sensitive phenotype (Fig. **S3A-B**), likely by counteracting the impact of darkness on cellular energy levels and hence offsetting the need to induce an adequate starvation response (Baena-González *et al*., 2007). Supporting this view, *hos1* showed no symptoms of low-energy stress under control conditions, where photosynthesis and respiration are not limited (Fig. **1**, day 0). Responses to low-energy conditions are under control of the SnRK1 signalling pathway, whose main components are the SnRK1 protein kinase and the C- and S-subclass of bZIP TFs (Baena-González *et al*., 2007; Mair *et al*., 2015; Pedrotti *et al*., 2018; Dröge-Laser and Weiste, 2018). Although we did not test specifically for defects in bZIP TFs, our data suggests that the defect of the *hos1* mutants in low-energy signalling relates at least in part to the SnRK1 kinase itself. Firstly, we show that HOS1 promotes tolerance to low-energy conditions (Fig. **1**, Fig. **S1** and Fig. **S2**). Secondly, we show that it interacts physically with SnRK1α1 (Fig. **2**) and that *hos1* has reduced accumulation of nuclear SnRK1α1 (Fig. **3**). Finally, we show that promoting the nuclear accumulation of SnRK1α1 in *hos1* by an NLS-fusion rescues to a large extent the defects of the *hos1* mutant in low-energy responses (Fig. **5**).

HOS1 has an E3 ubiquitin ligase activity that was shown to promote proteasome-dependent degradation of the ICE1 (Dong *et al*., 2006) and CO (Lazaro *et al*., 2012) transcriptional regulators. However, SnRK1 protein levels were not increased in *hos1* when analyzed from total protein extracts (Fig. **S4A-B**) or cytoplasmic and nuclear fractions (Fig. **3A-D**), indicating a non-proteolytic role of HOS1 in SnRK1 regulation. This is in accordance with reports of other HOS1 interactors that are regulated by HOS1 independently of its E3 ubiquitin ligase function (Lee *et al*., 2012; Jung *et al*., 2013; Kim, Lee, Jung, *et al*., 2017; Han *et al*., 2020). HOS1 is found in most land plants but the RING finger domain is not always present, lacking for example in the HOS1 proteins of moss (*Physcomitrium patens*), common bean (*Phaseolus vulgaris*) and soybean (*Glycine maxima*) (Jung *et al*., 2014). In contrast, the NUP ELYS motif is ubiquitous and is highly conserved, suggesting that non-proteolytic roles of HOS1, such as mRNA export and chromatin remodelling, could be more important than its E3 ubiquitin ligase activity (Jung *et al*., 2014; MacGregor *et al*., 2013). In this context, the fact that NLS-SnRK1α1 was able to rescue to a large extent the defects of *hos1* in low-energy responses (Fig. **5C-E**) suggests that at least one of the causes for these defects is the depletion of SnRK1α1 from the nucleus, and that HOS1 is important for the nuclear accumulation of SnRK1α1. This interpretation is further supported by the molecular and plant phenotypes of the NPC mutant *nup160* in response to prolonged darkness (Fig. **4**), which are similar to those of *hos1*. Given that NUP160 is required for anchoring HOS1 to the NPC (Li *et al*., 2020), this indicates that SnRK1 signalling requires HOS1 localization at the NPC and/or an adequate NPC function. Supporting this, NUP160 was also recently identified as a SnRK1α1 interactor by TurboID-based proximity labelling (van Leene *et al*., 2022), and the double *nup160/hos1* mutant showed the same molecular and plant phenotypes in response to prolonged darkness as its parents (Fig. **S5**). Altogether, this suggests that the HOS1 and NUP160 nucleoporins may act in concert to promote tolerance to low-energy stress *via* the SnRK1 kinase. Loss of HOS1 and/ or NUP160 may impact the function and structural integrity of the NPC (Lüdke *et al*., 2021), and thereby cause reduced accumulation of SnRK1α1 in the nucleus of the respective mutant plants. The fact that the nucleoporin NUP96 protein levels are reduced in *hos1* (Cheng *et al*., 2020) suggests that the structural integrity of the NPC may indeed be compromised in this mutant. Furthermore, several NUPs, including NUP160, were shown to be important for the nucleo-cytoplasmic distribution of key regulators of auxin, salicylic acid signalling and defense signalling components (Parry *et al*., 2006; Cheng *et al*., 2009; Du *et al*., 2016), and HOS1 was recently implicated in the nuclear localization of the PIF4 transcription factor (Zhang *et al*., 2020). Depletion of the NUP ELYS, with a region of homology with HOS1, in human cells resulted in lower NPC density and reduced nuclear size (Jevtić *et al*., 2019), but normal nuclear size could be restored by overexpression of importin α, arguing that the impact of ELYS on nuclear size is import-mediated.

How precisely HOS1 interferes with the nuclear accumulation of SnRK1α1 awaits further mechanistic clarification. The SnRK1 regulator PLEIOTROPIC REGULATORY LOCUS 1 [PRL1; (Bhalerao *et al*., 1999)] interacts with IMPORTIN ALPHA 3 [IMPA3/MOS6; (Németh *et al*., 1998)], raising the possibility that SnRK1 nuclear import is modulated by the PRL1/IMPA3 interplay. It is possible that the physical interaction detected between HOS1 and SnRK1α1 (Fig. **2A-B**) could occur in the context of SnRK1α1 transport to the nucleus. The fact that the nuclear accumulation of PIF4 is compromised in the *hos1* mutant (Zhang *et al*., 2020) and that HOS1 interacts with PIF4 (Kim, Lee, Jung, *et al*., 2017), supports the view that HOS1 may contribute to the regulation of nuclear protein accumulation. It further suggests that this may occur through interaction with specific cargo proteins, such as PIF4 or SnRK1α1.

The reduced nuclear accumulation of SnRK1α1 in *hos1* causes defective induction of starvation genes and increased susceptibility to prolonged darkness, consistent with reports that nuclear SnRK1α1 is required for transcriptional responses and survival under low-energy conditions (Cho *et al*., 2012; Ramon *et al*., 2019). Like mammalian AMPK, which translocates to the nucleus in response to several stress and hormonal stimuli (Kim *et al*., 2014; McGee *et al*., 2003; Kodiha *et al*., 2007; Suzuki *et al*., 2007), nuclear levels of SnRK1α1 were reported to increase under energy stress (Ramon et al., 2019), although such increase appeared to be specific to particular cell types (Blanco *et al*., 2019). In the case of *hos1*, nuclear accumulation of SnRK1α1 was constitutively reduced compared to Col-0, both in light (Fig. **3A-B**) and dark stress conditions (Fig. **3C-D**). Similarly, the differences in the nuclear accumulation of IAA17 and PIF4 between Col-0 and *nup* mutants are constitutive, being quite prominent under control conditions. Although the differences are enhanced by heat stress, this enhancement is proportionally small when compared to the basal situation (Zhang *et al*., 2020). On the other hand, the fact that in *hos1* the reduced accumulation of SnRK1α1 in the nucleus is not accompanied by a proportional increase in the cytoplasmic fraction is to be expected, given that the nucleus constitutes a small percentage (~8%) of the total cell volume (Huber and Gerace, 2007). Therefore, the SnRK1α1 depleted from the *hos1* nucleus is expected to contribute little to the much larger cytoplasmic fraction, remaining undetectable.

Given the involvement of HOS1 in processes that are highly interconnected, it is not always possible to identify the primary defect caused by altered HOS1 function. SnRK1 has likewise been associated to a wide variety of processes, some of which, including hypocotyl elongation (Simon, Kusakina, *et al*., 2018; Simon, Sawkins, *et al*., 2018), circadian function (Shin *et al*., 2017; Frank *et al*., 2018) and flowering time (Tsai and Gazzarrini, 2012; Jeong *et al*., 2015; Sanagi *et al*., 2021; Baena-González *et al*., 2007; Williams *et al*., 2014), are also affected by HOS1 (Kim, Lee, Jung, *et al*., 2017; MacGregor *et al*., 2013; Lazaro *et al*., 2012). A negative role in flowering regulation has been assigned to HOS1 and SnRK1α1, as shown by the early flowering phenotype of *hos1* (Lazaro *et al*., 2012) and by the delayed flowering of *SnRK1α1* overexpressor plants (Williams *et al*., 2014; Baena-González *et al*., 2007; Tsai and Gazzarrini, 2012). However, only the defects of *hos1* in the low-energy response but not in flowering could be rescued by increasing the presence of SnRK1α1 in the nucleus (Fig. **S6**), showing that nuclear depletion of SnRK1α1 does not account for all the *hos1* phenotypes.

It remains to be addressed whether the interaction between HOS1 and SnRK1 is of relevance for other processes e.g. circadian clock function, where opposing roles have been reported for these factors (Shin *et al*., 2017; Frank *et al*., 2018).

In summary, we show that depletion of HOS1 leads to defective responses to low-energy stress that can be traced back to a constitutive reduction of SnRK1α1 levels in the nucleus (Fig. **S7**). Altogether, our results demonstrate that the NPC component HOS1 is important for adequate SnRK1 signalling and thereby for plant tolerance to low-energy conditions.

## MATERIALS AND METHODS

All primers used in this study are provided in Supporting Information Table **S1**.

### Plant material and growth conditions

Arabidopsis seeds were vapor-sterilized and stratified at 4 °C for 2 d before sowing. Unless otherwise specified, plants were grown in soil under a 12 h light (100 μmol m^-2^ s^-1^), 22 °C/12 h dark, 18 °C regime. All *Arabidopsis thaliana* plants used in this study are in the Columbia (Col-0) background. The *hos1* [*hos1-3;* SALK_069312; *hos1-4;* EMS mutant, (Lazaro *et al*., 2012)], *nup160 [nup160-4* or *sar1-4;* SALK_126801; (Parry *et al*., 2006)], *NLS-SnRK1α1* and *βMYR-SnRK1α1* (Ramon *et al*., 2019), and *Snrk1α1* [*snrk1α1-3;* GABI_579E09; (Mair *et al*., 2015; Crozet *et al*., 2016)] lines have been previously described. The *hos1* mutant was crossed with *Snrk1α1, nup160, NLS-SnRK1α1* or *βMYR-SnRK1α1*, to obtain the respective multiple mutant plants. To generate a *HOS1* complementation line (*35S::HOS1-GFP/hos1*), the coding sequence of *HOS1* (At2g39810.1) was amplified from leaf cDNA using the primers indicated in Table **S1** and cloned into a pCB302-derived minibinary expression vector with a C-terminal GFP tag and under the control of the *35S* promoter (Baena-González *et al*., 2007). The construct was introduced into *Agrobacterium tumefaciens* (GV3101) and *hos1* plants were transformed by the floral dip method (Clough and Bent, 1998). BASTA^®^-resistant transformants were selected based on their segregation ratio (T2) and homozygosity (T3).

In plate assays, for experiments involving sugar supply or when phenotyping the *nup160/hos1* mutant, sterilized seeds were plated in 0.5 x Murashige and Skoog Basal Medium (MS) (pH 5.7) with or without 1% sucrose as indicated, and stratified at 4 °C for 2 d. Plates were then transferred to the same equinoctial conditions as plants grown in soil (see above) for 12 d, prior to the dark stress treatments. Survival was scored at 7+ 7 d, as the ability to retain chlorophyll in the shoot apex (as an indication of meristem viability) and the overall capacity to increase plant size and prevent desiccation.

### Dark stress treatments

For plant material and growth conditions prior to treatment, refer to that specific section. **Short dark treatment (3 h).** Fully expanded leaves of 4-week-old plants were detached from the rosettes at ZT6 and incubated on sterile MilliQ water in Petri dishes (3-4 detached leaves, from a minimum of 2 rosettes) for 3 h under light (L) (control; 100 μmol m^-2^ s^-1^) or darkness (D, covered with aluminum foil). L and D samples were collected at ZT9 for gene expression analysis. **Extended night treatment (9 h).** Two minutes before the start of the light period, 4-week-old Arabidopsis plants were either placed to a dark chamber at 22 °C (constant temperature) or left under the regular conditions as controls. Rosettes were collected at ZT9 after 9 h of extended darkness (D) or control light treatment (L) for protein extraction and Western-blot analysis. **Long dark treatment (2 d).** 4-week-old Arabidopsis plants (day 0) were placed at constant darkness and temperature (22 °C) for 2 d (day 2). Rosettes collected in each time point were used for gene expression analyses. **Long dark treatment (7 d).** Four-week-old Arabidopsis plants or 12 d-old seedlings (plate assays) were collected (day 0 samples) or placed at constant darkness and temperature (22 °C) for 7 d (day 7). Plants were then transferred back to the original conditions for recovery for 7 additional days (day 7+ 7). Rosettes collected in each time point were used for image acquisition and for measuring rosette fresh weight.

### Yeast two-hybrid (Y2H) assays

The full□length coding sequence of HOS1 (At2g39810.1) was cloned into pGADT7 in fusion with the GAL4 activation domain (AD). pGADT7 constructs were faced with pGBKT7 harboring full-length SnRK1α1 (At3g01090.1) fused to the DNA binding domain (BD) of GAL4. The empty vectors (EV) were used as negative controls. Yeast cells (Y2HGold) were transformed with both pGADT7 and pGBKT7 constructs and yeast growth was assessed in low-stringency media [-Leucine (L)/-Tryptophan(W)] for selection of co-transformants, and in high-stringency media [-L/-W/-Histidine (H)/-Adenine (A)] for selection of interactors.

### (Co)-Immunoprecipitation assays

Arabidopsis rosettes of 4-week-old plants (*35S::GFP* and *35S::HOS1-GFP/hos1*) were finely ground in liquid nitrogen. Proteins were extracted with immunoprecipitation (IP) buffer (1 ml/g fresh material) [50 mM Tris–HCl pH 8.0, 50 mM NaCl, 1 % (v/v) Igepal CA-630, 0.5 % (w/v) sodium deoxycholate, 0.1 % (w/V) SDS, 1 mM EDTA pH 8.0, 50 μM MG132, 50 mM N-ethylmaleimide, cOmplete protease inhibitor cocktail (1 tablet/20 mL, Roche 11697498001), and 1/500 (v/v) phosphatase inhibitor 2 (Sigma P5726) and phosphatase inhibitor 3 (Sigma P0044)]. After clearing samples by centrifugation (21130 *g*, 4 °C, 15 min), a small aliquot (30 μl) of each supernatant was taken and quantified using Pierce™ 660 nm assay reagent with the ionic detergent compatibility reagent. The remaining supernatant was incubated at 4 °C for 2 h under gentle rotation with 40 μl of μMACS anti-GFP MicroBeads (μMACS GFP Isolation Kit, Miltenyi, 130-091-125). Samples were thereafter loaded in μColumns (Miltenyi Biotec, 130-042-701) pre-equilibrated with 1 mL of IP buffer and allowed to flow through. Columns were washed four times with 200 μl of IP buffer and proteins eluted with 80 μl of elution buffer (Miltenyi, 130-091-125) at 95 °C. β-Mercaptoethanol (2 %) was added to eluates and supernatant samples prior to boiling for 5 min at 95 **°** C. Proteins were resolved by SDS-PAGE, wet-transferred to a PVDF membrane (110 V, 1 h at 4 **°** C), and immunodetected using α-SnRK1α1 and α-GFP antibodies.

### Protein Extraction

Arabidopsis rosettes of 4-week-old plants (Col-0, *hos1* and *HOS1-GFP/hos1* complementation lines) were finely ground using a TissueLyser II (Qiagen), and proteins were extracted using the IP buffer described in the “Immunoprecipitation assays” (above) (1,5 ml/g fresh material). After clearing samples by centrifugation (21130 *g*, 4 °C, 15 min), the supernatants were recovered and total protein was quantified using the Pierce™ 660 nm protein assay reagent with the ionic detergent compatibility reagent. Samples were denatured using 4x Laemmli solubilization buffer and boiling for 5 min at 95 °C. An equal protein amount per sample was loaded in the gels, samples were resolved by SDS-PAGE, wet-transferred to a PVDF membrane (110 V, 1 h at 4 °C), and immunodetected using antibodies against the indicated SnRK1 subunits or GFP (in the *GFP* and *HOS1-GFP/hos1* lines). Coomassie or Ponceau staining of the membranes was used as protein loading control.

### Nuclear Fractionation

Arabidopsis rosettes of 4-week-old plants were finely ground in liquid nitrogen and transferred to 2 ml tubes. The sample powder was added to 3x (w/v) of Nuclear Isolation Buffer (NIB) [20 mM Tris-HCl pH 8.0, 250 mM Sucrose, 5 mM MgCl_2_, 5 mM KCl, 5 mM EDTA pH 8.0, 14 mM β-Mercaptoethanol, 0.6 % (v/v) Triton X-100, 1 % (w/v) PVP40, Polyvinylpyrrolidone, cOmplete protease inhibitor cocktail (1 tablet/50 mL, Roche 11697498001), and 1/500 (v/v) phosphatase inhibitor 2 (Sigma P5726) and phosphatase inhibitor 3 (Sigma P0044)] and incubated under gentle rotation at 4 °C for 15 min. Samples were then filtered through two layers of Miracloth^®^ (Merck-Millipore; prewet with NIB) and centrifuged at 1240 *g*, 4 °C for 10 min. The supernatant, corresponding to the cytoplasmic fraction, was thereafter separated from the nuclei (pellet). Nuclei were washed four times with 2 ml of NIB, carefully resuspending the pellets with a brush and centrifuging at 1240 *g*, 4 °C for 10 min, each time. Nuclei were then resuspended in 200 μL of NIB (bis) [20 mM Tris-HCL pH 8.0, 250 mM Sucrose, 5 mM MgCl_2_, 5 mM KCl, 5 mM EDTA pH 8.0, cOmplete protease inhibitor cocktail (1 tablet/50 mL), and 1/500 (v/v) phosphatase inhibitor 2 and phosphatase inhibitor 3], added on top of 1 mL of a Percoll solution [15 % (v/v) Percoll, 20 mM Tris-HCL pH 8.0, 250 mM Sucrose, 5 mM MgCl_2_, 5 mM KCl, 5 mM EDTA pH 8.0, cOmplete protease inhibitor cocktail (1 tablet/50 mL), and 1/500 (v/v) phosphatase inhibitor 2 and phosphatase inhibitor 3] and centrifuged at 1240 *g*, 4 °C for 10 min. Supernatants were thereafter discarded and nuclei pellets were mixed with 20 μL of 4x Laemmli buffer, and boiled at 95 °C for 15 min, with brief vortexing every 5 min. Cytoplasmic fractions collected in the beginning were quantified using Pierce™ 660 nm assay reagent with the ionic detergent compatibility reagent. An equal protein amount of cytoplasmic samples, and whole nuclei samples were separated in a 15 % SDS-PAGE gel and analyzed with α-SnRK1α1, α-Histone H3, and α-Rubisco small subunit (RbcS) antibodies by Western-blotting.

### Antibodies and Western-blotting (WB)

The SnRK1α1 (1/4000) antibody was previously described (Belda-Palazón *et al*., 2020). The SnRK1β1 (1/500, anti-AKINB1, AS09460) was purchased from Agrisera. Phospho-SnRK1α1/2 (T175/176) was detected with an anti–phosphoT172-AMPKα antibody (1/1000 in 5 % BSA-TBS-Tween, referred to as P-AMPK; #2535, Cell Signalling Technologies). Histone H3 (1/5000, ab1791, Abcam,) and RbcS (1/5000, anti-RbcS, AS07 259, Agrisera) were used to characterize the nuclear and cytoplasmic fractions, respectively. The anti-GFP (1/1000, 11814460001, Roche) antibody was used to detect the corresponding tagged proteins. For Western-blotting, all primary antibodies were diluted in 1 % non-fat milk in Tris-buffered saline (TBS) (unless otherwise stated) and incubated with the membrane under gentle shaking for 16 h at 4 °C. Chemiluminescent detection was performed after incubation with a 1/10000 dilution of Peroxidase AffiniPure goat anti-rabbit or anti-mouse IgG (H+L; Jackson ImmunoResearch) in 1 % non-fat milk in TBS for 1 h at room temperature, using the ECL Western Blotting Detection Reagent (GE Healthcare) and the SuperSignal West Femto Maximum Sensitivity Substrate (Thermo Scientific).

### Gene Expression analyses

For RT-qPCR quantification of gene expression, RNA was extracted from detached leaves and rosettes of Arabidopsis plants using TRIzol Reagent (Ambion, Life Technologies). RNA was thereafter treated with RNase-Free DNase (Promega) and reverse transcribed (1 μg) using SuperScript III Reverse Transcriptase (Invitrogen, ThermoFisher), according to the manufacturer’s instructions. RT-qPCR was performed using the iTaq Universal SYBR Green Supermix (cat. No. 1725124, Bio-Rad) according to the manufacturer’s instructions in a total volume of 10 μL. The PCR program comprised an initial denaturation for 30 s at 95 °C and amplification by 40 cycles of 15 s at 95 °C and 1 min at 60 °C and was ran in an ABI QuantStudio 7 Real Time PCR machine (Applied Biosystems). Expression values of *DIN6, DIN1* and *DRM2* genes were normalized to the average mean of *UBC21* and *SAND* expression (Czechowski *et al*., 2005), using the 2^-ΔCt^ method for relative quantification (Livak and Schmittgen, 2001).

### Flowering assay

The number of rosette leaves at flowering (with a stem height of approximately 1 cm) was assessed in several plants of the different genotypes analyzed, grown in soil, under a long-day regime [16 h light (100 μmol m^-2^ s^-1^), 23 °C/8 h dark, 18 °C].

### Statistical analyses

Box and whiskers plots, violin plots, and scatter plots were used to represent the experimental results. In box and whiskers plots, lower and upper box boundaries represent the first and third quartiles, respectively, horizontal lines mark the median and whiskers mark the highest and lowest values. In scatter plots, bars represent the mean ± SEM. Dots represent individual datapoints. In the **RT-qPCR** plots, each data point refers to a whole rosette with the exception of Fig **S1A**, where it refers to a pool composed of 3-4 detached leaves from a minimum of 2 rosettes. Each experiment was treated as a batch within which the different genotypes were matched. Statistically significant differences between the different genotypes within each treatment or condition, were assessed by a two-way “randomized block” ANOVA, followed by a Tukey’s or a Sidak’s multiple comparisons test, with a 95% confidence interval. In the **fresh weight** plots, each data point refers to a whole rosette. For each genotype, individual fresh weight values were normalized to the average fresh weight at day 0 of the same genotype. Statistically significant differences between the different genotypes within each treatment or condition, were assessed by ordinary two-way ANOVA, with no matching or pairing selected, followed by a Tukey’s or a Sidak’s multiple comparisons test, with a 95% confidence interval. In the **protein quantification plots**, each data point refers to a normalized protein band. Each experiment was treated as a batch within which the different genotypes were matched. Statistically significant differences between the different genotypes were assessed by a ratio paired *t*-test, with a 95% confidence interval (two-tailed). In the **n° of rosette leaves** violin plots, each data point refers to a whole rosette. No matching or pairing was selected. Statistically significant differences between the different genotypes were assessed by ordinary one-way ANOVA, followed by a Tukey’s multiple comparisons test, with a 95% confidence interval. In the **% of survival** plots, each bar represents the mean ± SEM of the survival rates, for each genotype in each condition, from at least 8 independent assays. Each experiment was treated as a batch within which the different genotypes were matched, within each condition. Statistically significant differences between the different genotypes were assessed by either two-way ANOVA with Sidak’s multiple comparisons, or by ordinary one-way ANOVA, followed by a Tukey’s multiple comparisons test, with a 95% confidence interval. Plotting and statistical analyses were performed using the GraphPad Prism 9 version 9.1.1 for MacOS (GraphPad Software, LLC).

### Accession numbers

Sequence data from this article can be found in the Arabidopsis Genome Initiative database under the following accession numbers: *SnRK1α1*, At3g01090; *SnRK1α2*, At3g29160; *SnRK1β1*, At5g21170; *HOS1*, At2g39810; *NUP160*, At1g33410; *DIN6*, At3g47340; *DIN1*, At4g35770; *DRM2*, At2g33830; *UBC21*, At5g25760; *SAND*, At2g28390.

## ACKNOWLEDGEMENTS

We thank Vera Nunes from the IGC Model Organism Unit/Plant Facility for excellent plant management and Filip Rolland for kindly providing the *NLS-SnRK1α1* and *βMYR-SnRK1α1* lines. This work was supported by FCT - Fundação para a Ciência e a Tecnologia, I.P., through GREEN-IT - Bioresources for Sustainability R&D Unit (UIDB/04551/2020, UIDP/04551/2020) and LS4FUTURE Associated Laboratory (LA/P/0087/2020), PTDC/BIA-FBT/4942/2020, LISBOA-01-0145-FEDER-028128, PTDC/BIA-BID/32347/2017, EXPL/ASP-AGR/1329/2021, 2022.08339.PTDC, SFRH/BPD/116116/2016 (LM), PD/BD/114361/2016 (BP), and 2020.03177.CEECIND (EBG) and by the European Union Horizon 2020 research and innovation programme (Grant Agreement number: 867426 — ABA GrowthBalance — H2020-WF-2018-2020/H2020-WF-01-2018, awarded to BBP) 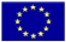

## AUTHOR CONTRIBUTIONS

L.M., A.C., and E.B.G. designed research; L.M., A.E., B.B.P., and B.P. performed the experiments; L.M. and A.E. analyzed data; E.B.G. supervised the research; E.B.G. and L.M. wrote the article with input from all coauthors.

---

**Supporting Fig. S1.**
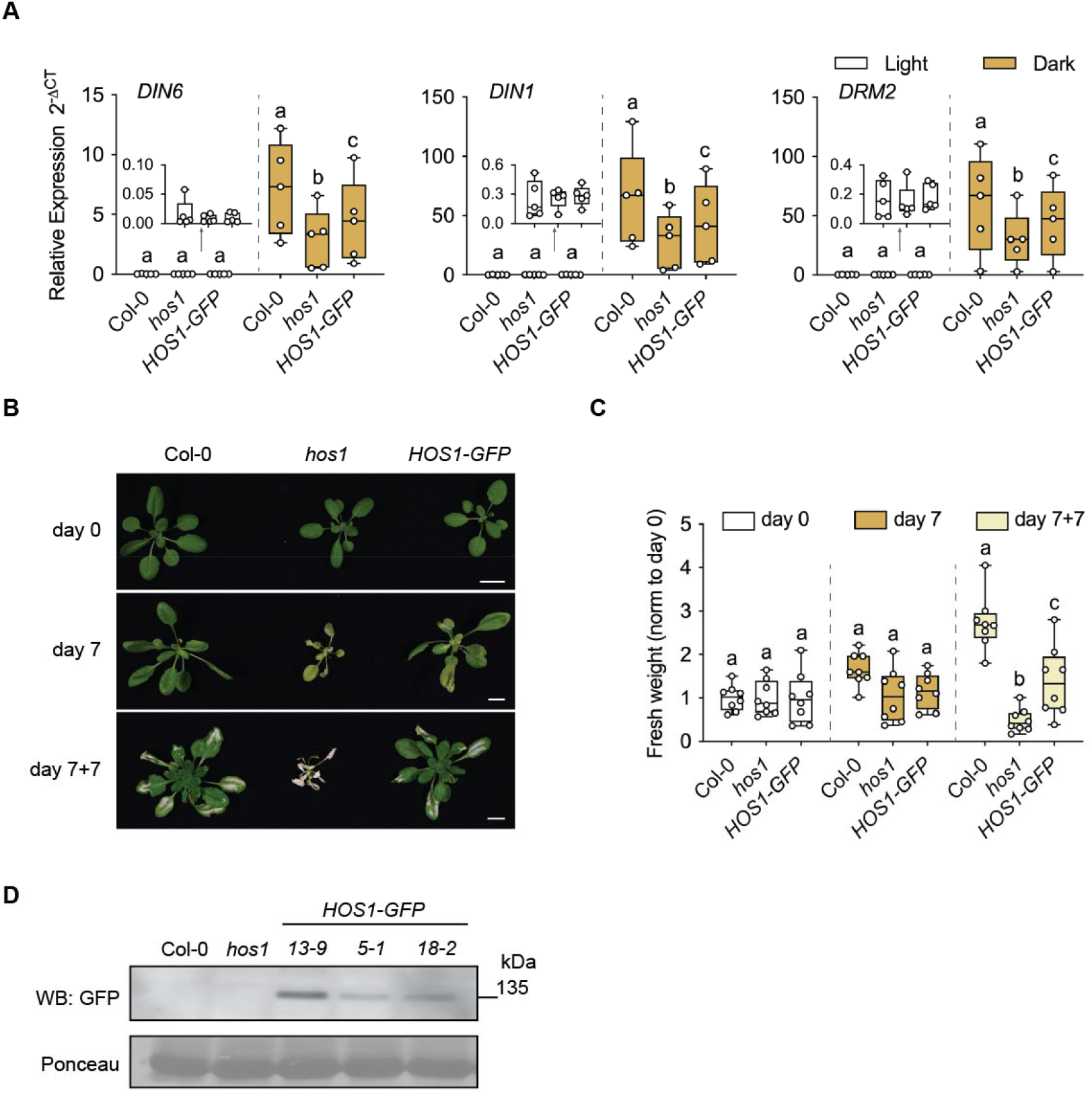
Tolerance of *hos1* and a *HOS1-GFP* complementation line to darkness. **A**. Expression of starvation genes (*DIN6, DIN1, DRM2*) measured by RT-qPCR from detached rosette leaves of 4-week-old Col-0, *hos1*, and *HOS1-GFP* plants (*HOS1-GFP/hos1* complementation line *13-9*, see below), after a 3 h light (L) or dark (D) treatment. Box and whiskers plots represent 5 biological replicates (each composed of 3-4 detached leaves from a minimum of 2 rosettes). **B.** Representative pictures of 4-week-old Col-0, *hos1*, and *HOS1-GFP* plants (day 0) placed in constant darkness for 7 d (day 7) and transferred back to the initial photoperiod for 7 additional days (day 7+7). Scale bar = 1 cm. **C.** Rosette fresh weight determined at days 0, 7 and 7+7, and normalized to day 0 for each genotype. Box and whiskers plot represents 8 plants per genotype and condition (grown in 3 independent batches). Different letters denote statistically significant differences between genotypes within each condition (L, D, or days 0, 7 and 7+7) as determined by two-way ANOVA with Tukey’s multiple comparisons test (p<0.05). **D.** HOS1-GFP protein levels in 4-week-old rosettes of Col-0, *hos1*, and 3 independent *HOS1-GFP/hos1* complementation lines (*13-9, 5-1* and *18-2*). Western-blot (WB) with an anti-GFP antibody and Ponceau staining as loading control.

**Supporting Figure S2.**
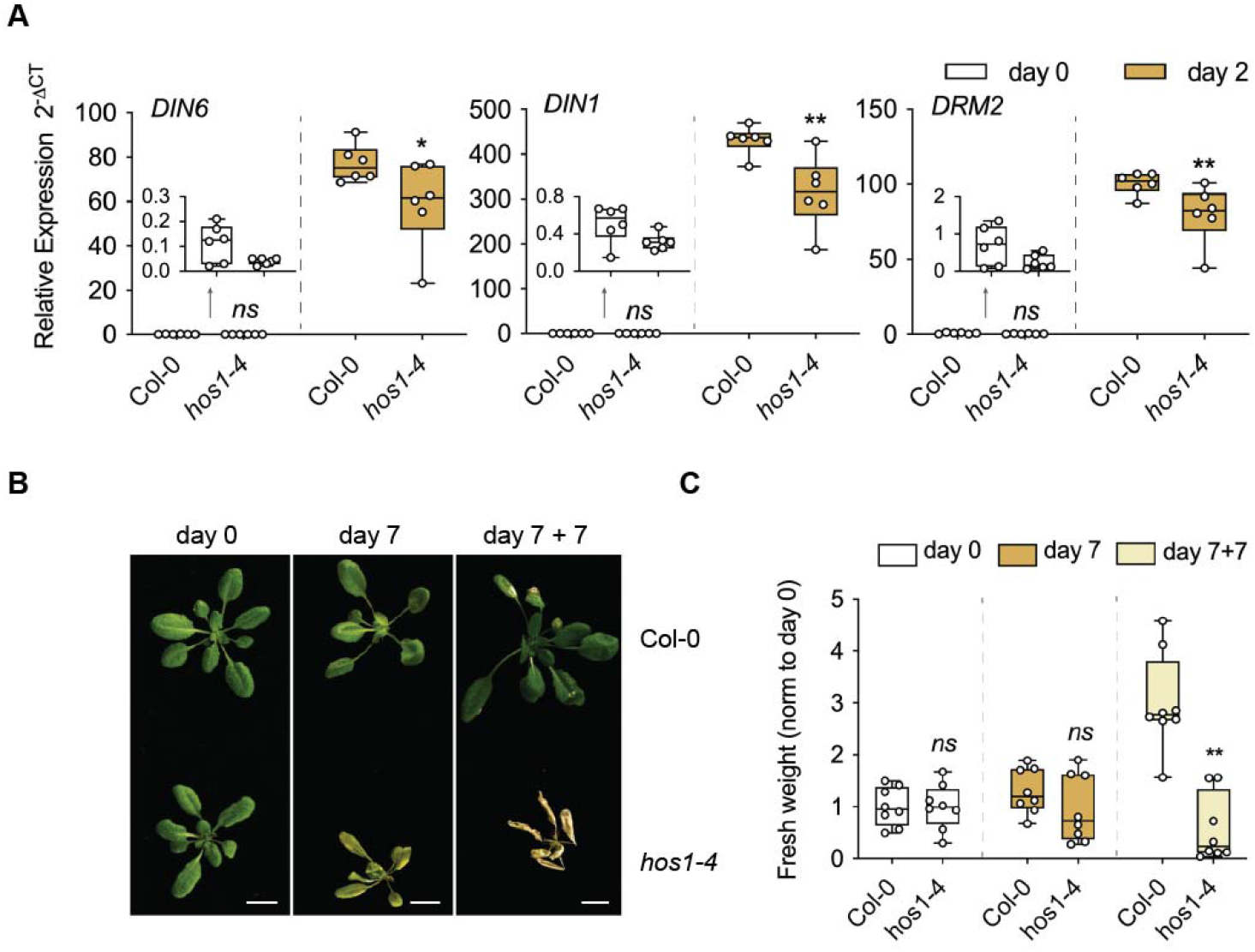
Tolerance of the *hos1-4* mutant to prolonged darkness. **A.** Induction of starvation genes (*DIN6, DIN1, DRM2*) measured by RT-qPCR from rosettes of 4-week-old Col-0 and *hos1-4* mutant plants collected at day 0 and after 2 d of dark treatment (day 2). Box and whiskers plots represent 6 biological replicates (each consisting of 1 rosette per genotype and condition, grown in 3 independent batches). **B.** Representative pictures of 4-week-old Col-0 and *nup160* plants (day 0) placed in constant darkness for 7 d (day 7) and transferred back to the initial photoperiod for 7 additional days (day 7+7). Scale bar = 1 cm. **C.** Rosette fresh weight determined at days 0, 7 and 7+7, and normalized to day 0 for each genotype. Box and whiskers plots represent 8 plants per genotype and condition (grown as 4 independent batches). Asterisks denote statistically significant differences between Col-0 and *hos1-4* within each condition (days 0, 2, 7 and 7+7), as determined by two-way ANOVA with Sidak’s multiple comparisons test. (*) p< 0.05; (**) p< 0.01; (*ns*), non-significant.

**Supporting Figure S3.**
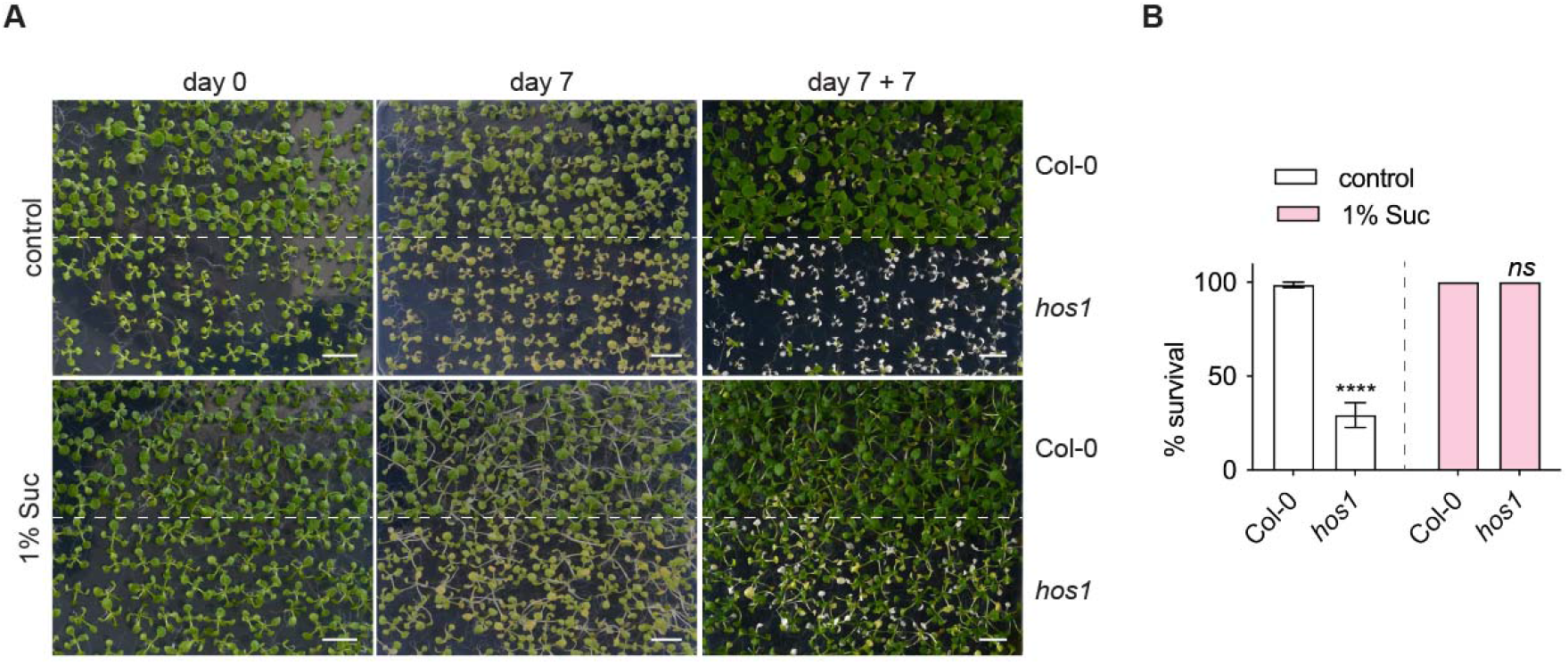
Tolerance of *hos1* to prolonged darkness in the presence of sugar. **A.** Representative images of 12 d-old seedlings (day 0) of Col-0 and *hos1* grown in 0.5 MS (control) or 0.5 MS 1% sucrose (1% Suc) plates, placed in constant darkness for 7 d (day 7) and transferred back to the initial photoperiod for 7 additional days (day 7+7). **B.** Quantification of survival at day 7+7, for each genotype and condition. Bars represent the mean percentage survival (%) ± SEM of a minimum of 8 independent batches (with a total amount of at least 350 seedlings per genotype and condition). Asterisks denote statistically significant differences between Col-0 and *hos1*, as determined by two-way ANOVA with Sidak’s multiple comparisons test. (****) p< 0.0001; (*ns*), non-significant.

**Supporting Fig. S4.**
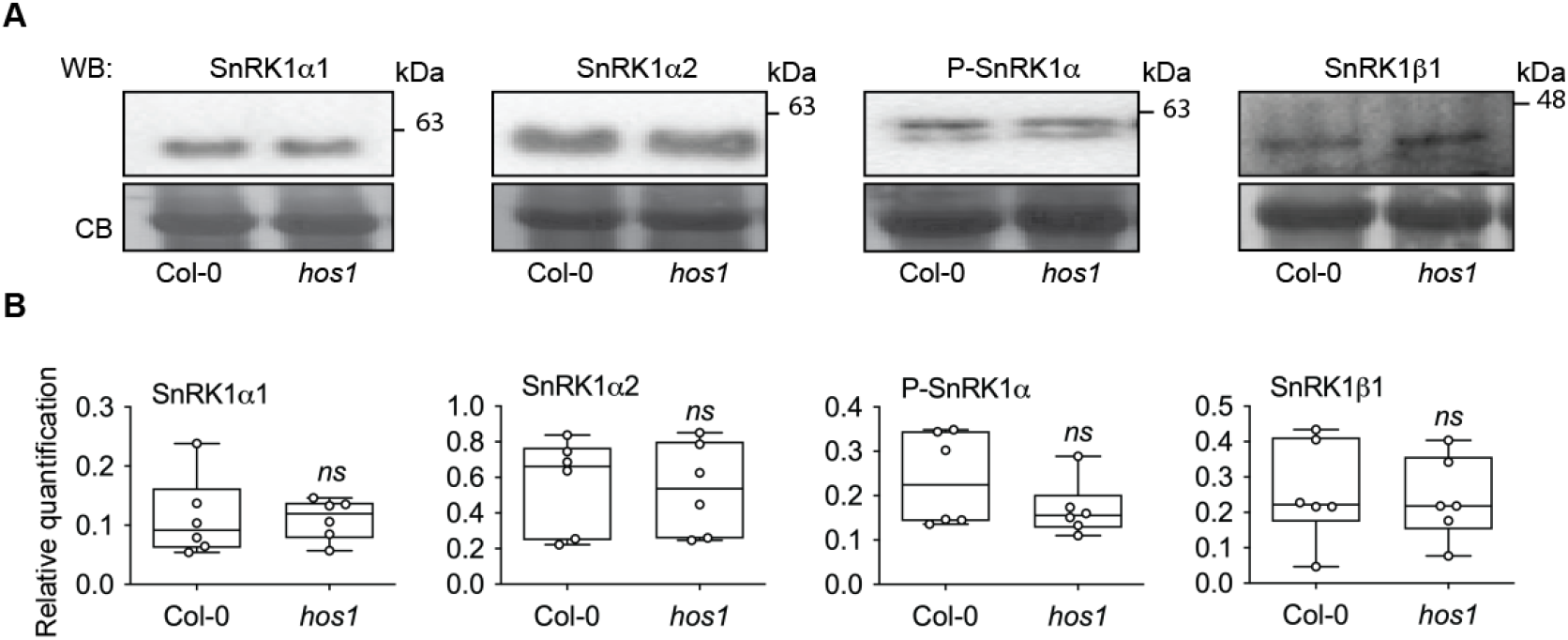
Accumulation of SnRK1 subunits in *hos1*. **A.** Levels of SnRK1 proteins in rosettes of 3-week-old Col-0 and *hos1* plants. Representative Western-blots (WB) with antibodies against several SnRK1 subunits (SnRK1α1, SnRK1α2, and SnRK1β1) and the T-loop phosphorylation of SnRK1α1 and SnRK1α2 (P-SnRK1α). Coomassie blue (CB) staining as loading control. **B.** Quantification of SnRK1 protein levels relative to the CB staining. Box and whiskers plots represent 6 biological replicates (each consisting of 1 rosette per genotype, grown in 6 independent batches). (*ns*) denotes no statistically significant differences between Col-0 and *hos1*, as determined by a ratio paired *t*-test.

**Supporting Fig. S5.**
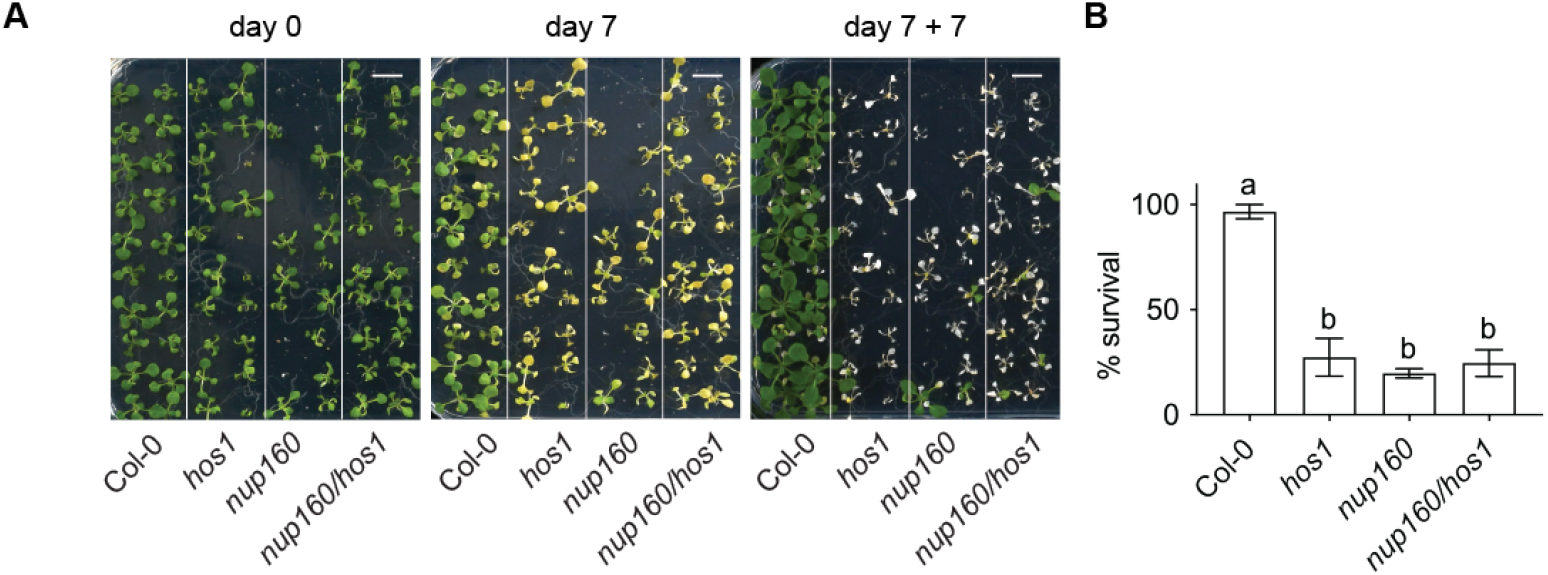
Tolerance of the NPC mutants *hos1* and *nup160* to prolonged darkness. **A.** Representative images of 12-day-old seedlings (day 0) of Col-0, *hos1, nup160* and *nup160/hos1* grown in 0.5 MS plates, placed in constant darkness for 7 d (day 7) and transferred back to the initial photoperiod for 7 additional days (day 7+7). **B.** Quantification of survival at day 7+7 for each genotype. Bars represent the mean percentage survival (%) ± SEM of a minimum of 5 independent batches (with a total amount of at least 65 seedlings per genotype). Different letters denote statistically significant differences between genotypes, as determined by one-way ANOVA with Tukey’s multiple comparisons test (p<0.05).

**Supporting Fig. S6.**
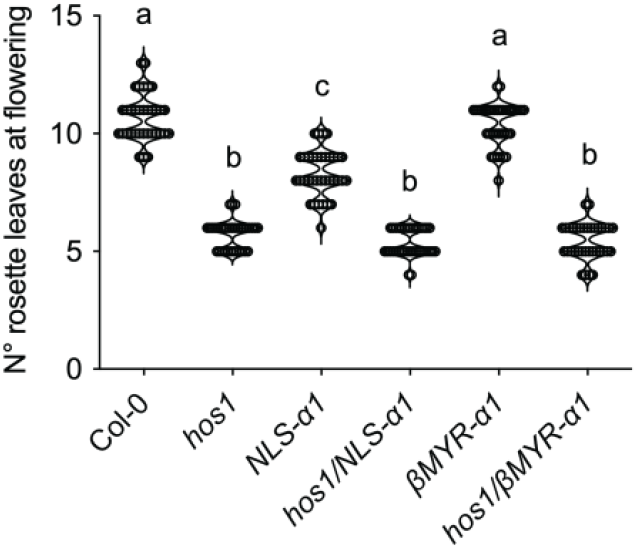
Impact of an *NLS-SnRK1α1* transgene on the flowering phenotype of *hos1*. Col-0, *hos1*, and *NLS-SnRK1α1* (*NLS-α1*) or *βMYR-SnRK1α1* (*βMYR-α1*) alone or crossed with *hos1*, grown in long-day conditions (16:8 h; 23:18 °C). Number of rosette leaves in each plant at the time of flowering. Violin plot represents 34 plants per genotype (grown in 4 independent batches). Different letters denote statistically significant differences between genotypes, as determined by one-way ANOVA with Tukey’s multiple comparisons test (p<0.05).

**Supporting Figure S7.**
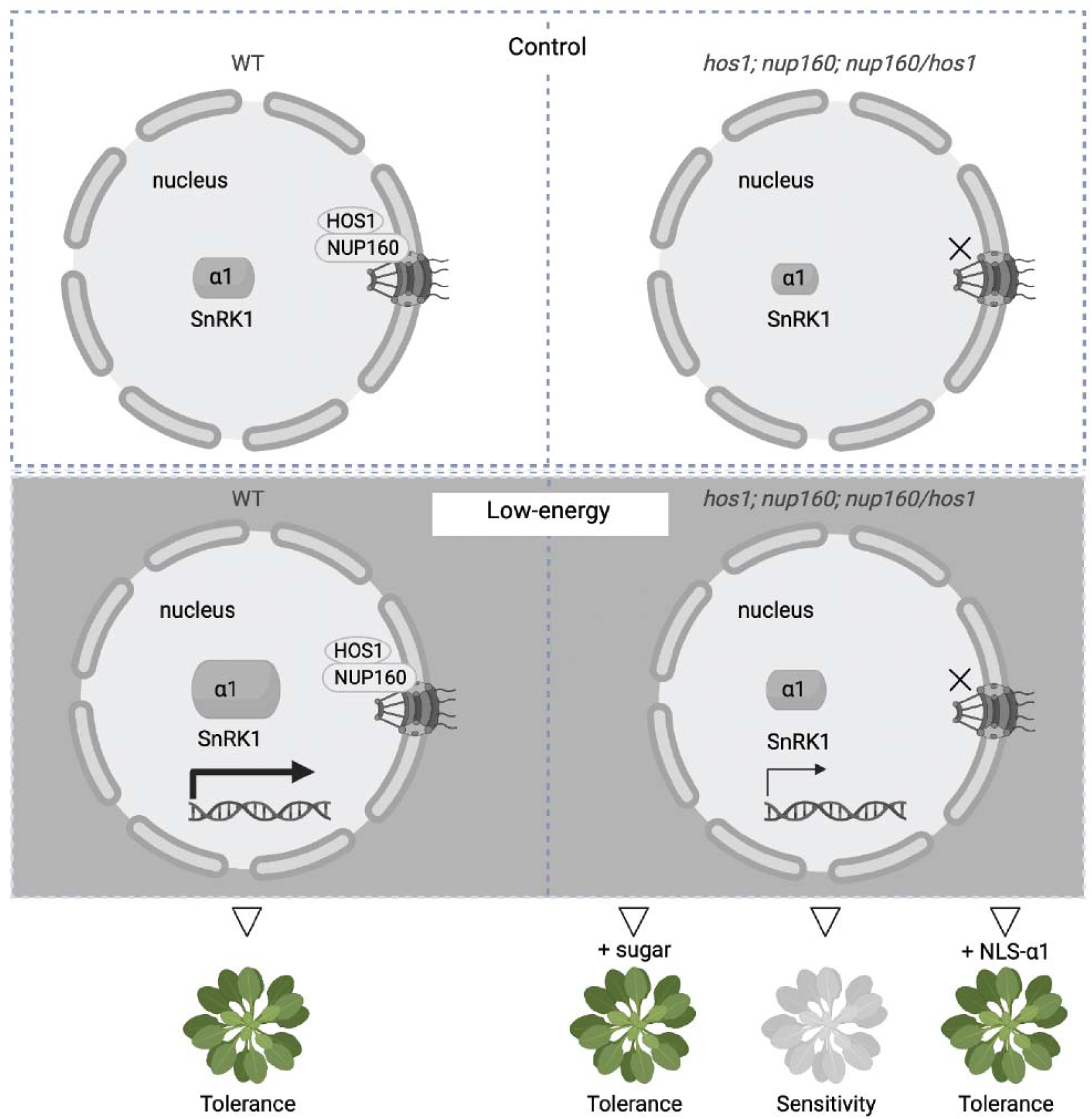
Model for the role of HOS1 in low-energy responses. In plants, low-energy conditions can derive from environmental stresses that impair photosynthesis or respiration. Plant tolerance to low-energy relies on a central energy sensor and stress regulator, the SnRK1 protein kinase. The presence of the SnRK1 catalytic subunit (SnRK1α1) in the nucleus is essential for the transcriptional reprogramming needed to balance energy levels (e.g. induction of “starvation” genes). Depletion of the nuclear pore complex (NPC) component HOS1 causes constitutive defects in the nuclear accumulation of SnRK1α1. Accordingly, knockout mutants of HOS1 and NUP160, a nucleoporin that anchors HOS1 to the NPC, show compromised responses to low-energy stress. These defects can be largely reverted by sugar supplementation and through nuclear mobilization of SnRK1α1 *via* an *NLS-α1* transgene. Created with BioRender.com.

**Supporting Table 1.**
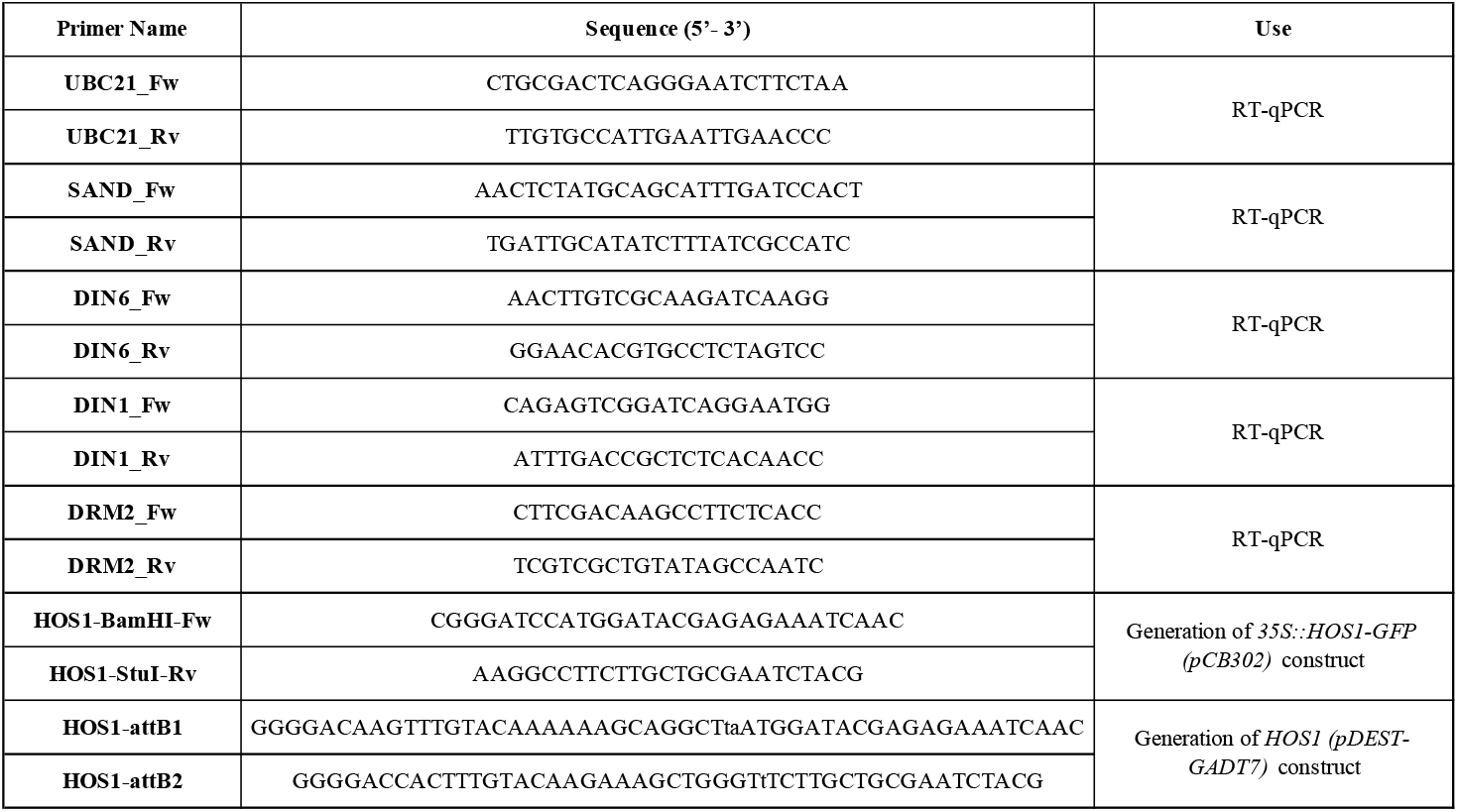
List of primers used.

